# AAV-mediated expression of mouse or human GLDC normalises metabolic biomarkers in a GLDC-deficient mouse model of Non-Ketotic Hyperglycinemia

**DOI:** 10.1101/2023.12.15.571844

**Authors:** Kit-Yi Leung, Chloe Santos, Sandra C.P. De Castro, Diana Gold Diaz, Andrew J. Copp, Simon Waddington, Nicholas DE Greene

## Abstract

Non-Ketotic Hyperglycinemia (NKH) is a rare inborn error of metabolism caused by impaired function of the glycine cleavage system (GCS) and characterised by accumulation of glycine in body fluids and tissues. NKH is an autosomal recessive condition and the majority of affected individuals carry mutations in *GLDC* (glycine decarboxylase). Current treatments for NKH are not effective or curative. As a monogenic condition with known genetic causation, NKH is potentially amenable to gene therapy. An AAV9-based expression vector was designed to target sites of GCS activity. Using an ubiquitous promoter to drive expression of a GFP reporter, transduction of liver and brain was confirmed following intra-venous and/or intra-cerebroventricular administration to neonatal mice. Using the same capsid and promoter with transgenes to express mouse or human GLDC, vectors were then tested in a GLDC-deficient mice that provide a model of NKH. GLDC-deficient mice exhibited elevated plasma glycine concentration and accumulation of glycine in liver and brain tissues as previously observed. Moreover, the folate profile indicated suppression of folate one carbon metabolism (FOCM) in brain tissue, as found at embryonic stages, and reduced abundance of FOCM metabolites including betaine and choline. Neonatal administration of vector achieved reinstatement of *GLDC* mRNA and protein expression in GLDC*-*deficient mice. Treated GLDC*-*deficient mice showed significant lowering of plasma glycine, confirming functionality of vector expressed protein. AAV9-GLDC treatment also led to lowering of brain tissue glycine, and normalisation of the folate profile indicating restoration of glycine-derived one-carbon supply. These findings support the hypothesis that AAV-mediated gene therapy may offer potential in treatment of NKH.

## 1. Introduction

Non-Ketotic Hyperglycinemia (NKH; glycine encephalopathy) is a life-limiting autosomal recessive disease, characterised by accumulation of glycine in body fluids and tissues [1–3]. Severe NKH presents in neonates, with lethargy, respiratory problems (apnea) that may require assisted breathing, hypotonia and seizures. Myoclonic jerks and burst suppression pattern on EEG are frequently present in neonates but the type, frequency and severity of seizures change in infants, to include hypsarrhythmia, burst suppression and multifocal epilepsy [4]. This clinical course highlights opportunity for early therapy that could prevent secondary epilepsy-related neurological damage. Surviving infants suffer profound neurological impairment with complex epilepsy and severe psychomotor retardation [1,5]. Around one-third of affected individuals die within the first year and the median age of death is 8 years.

NKH is caused by loss of function of the glycine cleavage system (GCS), an evolutionarily conserved mitochondrial protein complex that acts to decarboxylate glycine with concomitant transfer of a one-carbon group to tetrahydrofolate (THF) in folate one-carbon metabolism (FOCM)[2,6]. The GCS has four protein components, of which glycine decarboxylase (GLDC) and aminomethyltransferase (AMT) are thought to be GCS-specific, whereas GCS H protein (GSCH) and dihydrolipoamide dehydrogenase (DLD) have additional roles in other enzyme complexes [7,8]. Approximately 80% of individuals with NKH carry mutations in *GLDC,* with the majority of the other cases being attributable to mutations in *AMT* [6,9], and a small number to *GCSH* [10]. In a subset of cases, described as ‘variant NKH’ mutations in GCS-encoding genes are not detected and, in some individuals, mutations have been identified in the enzyme pathway responsible for synthesis of the lipoate co-factor, which is responsible for lipoylation of GCSH and other proteins [11,12].

In addition to severe NKH that presents in neonates or infants, it has become evident that a wider spectrum of disease is associated with GCS mutations. Attenuated NKH appears to result from *GLDC* or *AMT* mutations that result in partial loss of protein function, with heterogeneity in age of onset and a less severe neurological outcome [5], but still a debilitating disease. Identification of *GLDC* mutations in children or adults with mental retardation and behavioural problems [13–15], further suggests that mild or late-onset NKH may go unrecognised in some individuals.

Current treatment for NKH is largely focussed on control of glycine and amelioration of the predicted epileptogenic effects of glycine. Administration of sodium benzoate can lower glycine levels via stimulation of the glycine conjugation pathway in liver and kidneys, resulting in the production of benzoylglycine (hippurate) that is excreted in urine [16,17]. In addition, multiple anti-epileptics and the NMDA-receptor antagonist dextromethorphan are frequently used in order to mitigate the effects of excess glycine in the brain [18–20]. These treatments are not curative, and do not appreciably improve neurodevelopment, while benzoate treatment has associated side-effects and requires co-administration of antacids [17],[21].

While current treatments for NKH are not effective, the continuum of clinical severity from neonatal onset, severe NKH to milder and later-onset forms suggests that therapies which reinstate GCS function may have a beneficial effect. Unlike current treatments, restoration of GCS function would achieve both lowering of glycine and supply of glycine-derived one-carbon groups to FOCM. Gene therapy vectors provide potential to deliver a *GLDC* transgene to endogenous sites of expression in the liver and brain, and this approach would potentially be applicable to any loss of function mutation in *GLDC*.

Adeno-associated virus (AAV) based vectors are based on the AAV genome with replacement of the *rep* and *cap* genes by the therapeutic transgene and promoter. AAV vectors enter the target cell nucleus and exist primarily in episomal form with rare integration into the host chromosomes. Hence, long-term expression of transgenes may be achieved in non-dividing cells, such as neurons, although the vector may be ‘diluted’ in tissues undergoing cell proliferation [22,23]. The cell-type specificity (tropism) of AAV is determined by the capsid serotype. AAV9 has been found to be effective in crossing the blood brain barrier and can transduce neurons and non-neuronal cells, including astrocytes [22,24]. AAV9-based vectors have been used effectively in mouse models of several monogenic diseases including Krabbe disease, Niemann Pick type C and amyotrophic lateral sclerosis [25–30]. Successful testing of AAV9 vectors in animal models led to their implementation for clinical applications, for example, in treatment of spinal muscular atrophy [31–33].

As a model for testing potential novel treatments, mice that are homozygous for a gene-trap allele of *Gldc* (*Gldc^GT1^*) provide a GLDC-deficient model for NKH [34]. *Gldc^GT1/GT1^* mice exhibit 90% reduction in *Gldc* mRNA abundance, undetectable GCS activity and hallmark features of NKH, including elevated glycine concentrations in blood, urine and tissues [34]. In addition to glycine, GLDC-deficient mice display tissue-dependent accumulation of multiple glycine derivatives including acylglycines and guanidinoacetate, which is reversible by liver-specific rescue of *Gldc* expression [34,35]. In accordance with predicted effects of GCS loss of function [2], GLDC*-*deficient mice exhibit suppression of FOCM, with accumulation of tetrahydrofolate (THF) at the expense of folates that carry one-carbon groups [34,36]. In the current study we made use of the GLDC*-*deficient mouse model to test AAV9 vectors, owing to their ability to transduce liver and brain, the key target tissues in NKH.

## 2. Materials and Methods

### 2.1 *Gldc-*deficient mouse model of NKH

Animal studies were carried out under regulations of the Animals (Scientific Procedures) Act 1986 of the UK Government, and in accordance with the guidance issued by the Medical Research Council, UK in Responsibility in the Use of Animals for Medical Research (July 1993).

*Gldc*-deficient mice (*Gldc^GT1^*) carrying a gene-trap allele of *Gldc* were previously generated and characterised [34,35]. Mice used in the current study were on a C57BL/6J background. Experimental litters were generated by overnight timed mating of heterozygous (*Gldc^GT1/+^*) male and female mice. Pregnant females were supplemented with formate in the drinking water (30 mg/ml) from day 7 to 14 of pregnancy to prevent neural tube defects and hydrocephalus among the offspring [34]. Mice were genotyped by PCR of genomic DNA [34].

### 2.2 AAV vectors and administration

AAV9-CBh-eGFP was obtained from Vector Biolabs (Malvern, PA, USA). *GLDC* cDNA sequences were synthesised (GenScript) and cloned into pAAV-ITR plasmid (containing AAV2 ITRs) downstream of a hybrid chicken beta-actin (CBh) promoter. Plasmids and vector were verified by sequencing to confirm encoding of wild-type protein (mouse NP_613061, human NP_000161). Vector production was performed by Vector Biolabs (Malvern, PA, USA). pAAV plasmid was co-transfected with Ad Helper and AAV9 rep/cap vector into HEK293 cells and, following cell harvesting, vector was purified on two CsCl gradients to obtain titre of 2.1 x 10^13^ GC/ml for GLDC vectors and 3.9 x 10^13^ GC/ml for eGFP vector.

For neonatal administration, mice were genotyped from tail tip sample at P0 and injected with vector or vehicle (control) into the facial vein (intra-venous; 10 µl) and/or by intra-cerebroventricular route at P1. At P20, intra-venous administration of vector was achieved by injection of the tail vein (10 µl).

### 2.3 Collection of blood and tissues

Blood was collected by terminal cardiac exsanguination, transferred to lithium-heparin tubes (BD Microtainer) and immediately centrifuged for the isolation of plasma. Tissues were rinsed in PBS and snap frozen on dry ice to store at 80 C or fixed in 10% formalin overnight.

### 2.4 Immunohistochemistry

For histological analysis, formalin-fixed mouse brains were dehydrated, embedded in paraffin wax and sectioned at 7 µm thickness. For immunohistochemistry, primary and secondary antibodies were anti-GFP (chicken polyclonal, 1:300; Abcam), anti-GLDC (rabbit polyclonal, 1:500; Atlas Antibodies), Alexafluor goat anti-chicken (1:250; Invitrogen) and Alexafluor goat anti-rabbit (1:250, Invitrogen). For nuclear staining, cells were incubated with 4,6-diamidino-2-phenylindole (DAPI, 1:10,000 in PBS). Fluorescent images were collected on an Axiophot microscope (Zeiss) with a DC500 camera (Leica), using FireCam software (Leica).

### 2.5 Quantitative real time reverse transcription PCR

RNA was prepared using TRIzol reagent (Invitrogen), followed by chloroform extraction and RNA precipitation. First strand cDNA synthesis was performed using random hexamers (Superscript VILO cDNA synthesis kit). The abundance of *Gldc* mRNA was analysed by qRT– PCR (iTaq, BioRad) on BioRad CFX system, with each sample analysed in triplicate. Primers were located in exons 2 and 4 (5’ -AGCATTGATGAGCTCATCGAG-3’ and 5’ - TCCAGCAGGGAAGCGTTGGC-3’) of mouse *Gldc* and (5’-GCCCCTAGAAGCAGAGACTC-3’ and 5’-GGCACGGTTTTCTCGATCAG-3’) of human *GLDC* to amplify the wild-type but not the mutant transcript. Results were normalized to abundance of *Gapdh* expression.

### 2.6 Immunoassay and western blot and analysis

Brain and liver tissues were homogenised using handheld sonicator in RIPA buffer containing protease inhibitor cocktail and protein concentration in the lysate determined (Qubit). GLDC abundance was determined using anti-GLDC antibody (Atlas antibodies) and chemi-luminescent detection following separation on a capillary-based system (Jess, Bio-Techne) with normalisation to total protein. For western blot, protein samples (equal total loading) were electrophoresed on NUPAGE 10% gels and transferred onto PVDF membrane. GLDC protein was detected by anti-GLDC and GAPDH (detected using anti-GAPDH, Santa Cruz) was used as loading control. Following chemiluminescent detection, films were scanned using a GS800 densitometer (BioRad).

### 2.7 Glycine analysis

Tissue extracts were prepared by sonication in PBS containing 1x protease inhibitor cocktail (Roche). An aliquot was removed for protein determination by the Qubit assay. Tissue debris were removed by centrifugation (12,000 g at 4 C). Plasma and tissue lysate glycine were measured following the Kairos amino acid sample preparation and method (Waters corporation, UK). Briefly, protein was removed by the addition of 5-sulphosalicylic acid containing a specified quantity of amino acid internal standard in a ratio of 1:1 (sample:internal standard solution). Samples were derivatised using AccQ.Tag reagent. Glycine was resolved and detected using CORTECS UPLC C18, 1.6 μm, 2.1 x 150 mm column on an UPLC coupled to XEVO-TQS mass spectrometer (Waters Corporation, UK). Betaine and choline were analysed by LC-MS/MS (Metabolon Ltd, USA).

### 2.8 Folate analysis

Tissue extracts were prepared by homogenisation of samples by sonication in buffer containing 20 mM ammonia acetate, 0.1% ascorbic acid, 0.1% citric acid and 100 mM dithiothreitol at pH 7. An aliquot was removed for protein determination by the Qubit assay. Protein was removed by precipitation by addition of two volumes of acetonitrile and centrifugation (12,000 g at 4°C). Supernatants were transferred, lyophilized, stored at 80°C and re-suspended in dH2O before analysis. Folate analysis was carried out by ultraperformance liquid chromatography tandem mass spectrometry (UPLC-MS/MS) as described previously [36,37]. Lyophilized samples were resuspended in 30 ml water (milli-Q) and centrifuged for 5 min at 12,000 g at 4 C. Metabolites were resolved by reversed-phase UPLC (Acquity UPLC BEH C18 column, Waters Corporation, UK). Solvents for UPLC were as follows: Buffer A, 5% methanol, 95% Milli-Q water and 5 mM dimethylhexylamine at pH 8.0; Buffer B, 100% methanol, 5 mM dimethylhexylamine. The column was equilibrated with 95% Buffer A: 5% Buffer B. The sample injection volume was 25 ml. The UPLC protocol consisted of 95% Buffer A: 5% Buffer B for 1 min, followed by a gradient of 5–60% Buffer B over 9 min and then 100% Buffer B for 6 min before re-equilibration for 4 min. The metabolites were eluted at a flow rate of 500 nl/min and the wash step with 100% Buffer B was at flow rate of 600 nl/min. The UPLC was coupled to a XEVO-TQS mass spectrometer (Waters Corporation) operating in negative-ion mode using the following settings: capillary 2.5 kV, source temperature 150 C, desolvation temperature 600 C, cone gas flow rate 150 l/h, and desolvation gas flow rate 1,200 l/h. Folates were measured by multiple reaction monitoring with optimized cone voltage and collision energy for precursor and product ions [36,38].

### 2.9 Statistical Analysis

Pairwise comparisons were made by t-test. Multiple groups were compared by ANOVA with post-hoc analysis by Holm-Sidak method.

## 3. Results

### 3.1 AAV-mediated of GFP reporter following neonatal administration in mice

Among tissues in mammals and birds, GCS activity is highest in liver, kidney and brain, correlating with localisation of *GLDC* mRNA [2,39]. Liver activity of the GCS is sufficient to regulate circulating glycine levels in a non-tissue autonomous manner in mice [35]. However, GCS-mediated supply of glycine-derived one-carbon units in mitochondrial FOCM is predicted to be tissue autonomous. We therefore aimed to target liver and/or brain for AAV-mediated reinstatement of *Gldc* expression. The AAV9 capsid was selected based reported transduction of hepatocytes and CNS following administration to neonatal mice [40–42].

Although some studies have reported that vector genomes of up to 6 kb or even greater can be packaged in AAV vectors, the capacity is typically considered to be up to approximately 5 kb, corresponding to the size of the parental genome [43,44]. To drive broad expression in transduced tissues and to circumvent packaging limitations we designed a vector using AAV2 ITRs, hybrid chicken beta-actin (CBh) promoter and the GLDC-expressing transgene.

To assess transduction of target tissues and activity of the CBh promoter we performed preliminary studies using an eGFP transgene, packaged as AAV9-CBh-eGFP (Figure 1A). Vector was administered to wild-type (C57Bl/6) neonatal mice at post-natal day 1 (P1) by intravenous (iv) injection into the facial (temporal) vein, intracerebroventricular (ic) injection into the lateral ventricle, or by both iv and ic (iv + ic) route. A further group of mice were administered vector at P20 via iv injection into the tail vein. Mice were sacrificed 3 weeks after vector administration for analysis of GFP mRNA and protein expression.

**Figure 1.**
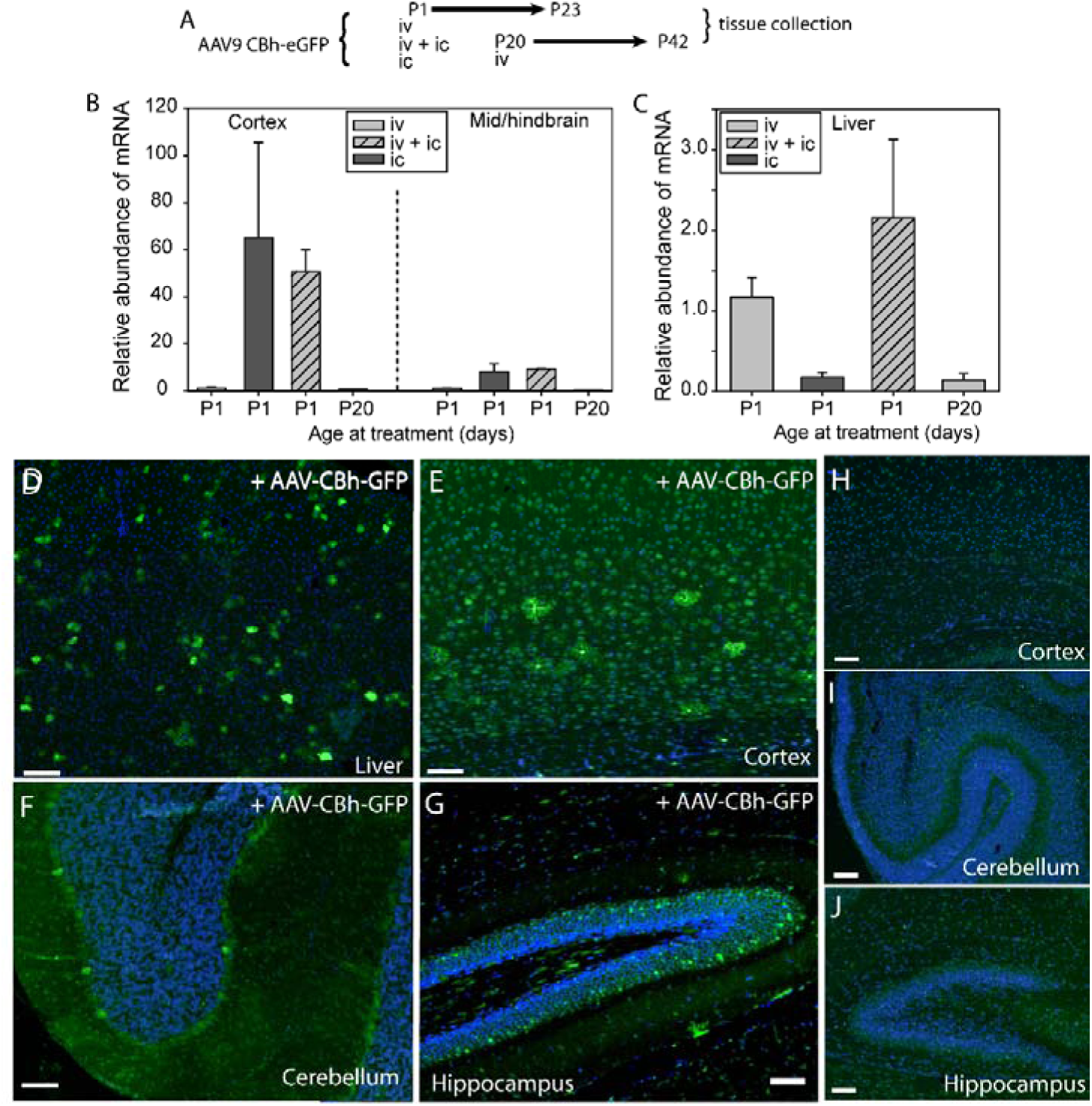
Evaluation of capsid and promoter combination for expression of transgene in moue brain. (A) AAV9-CBh-eGFP vector was administered to neonatal wild-type mice at post-natal day 1 (P1) ic, iv or iv and ic routes or at P20 by iv route. (B-C) *GFP* mRNA was detected in (B) brain and (C) liver. (D-G) Among mice treated with AA9-CBh-eGFP vector, GFP protein was detected by immunostaining in (D) liver (iv or iv +ic route at P1) and (E-G) different brain regions (ic or iv +ic route) including (E) cortex, (F) cerebellum, and (G) hippocampus. Examples are from mice treated by iv + ic route. (H-J) Low level of background fluorescence in samples from untreated mice. Scale bar represents 100 µm.

Expression of GFP mRNA was detected in both liver and brain, with varying abundance dependent on route and stage of administration. At 3 weeks post-treatment, expression in liver was higher in mice that were treated via iv or iv + ic routes at P2 compared with ic only, while expression following iv administration treatment at P20 was minimal compared with treatment at P2 (Figure 1B). Brain tissue was divided into anterior (olfactory bulbs and cortex) and posterior (mid and hindbrain) samples for mRNA quantification. Abundant GFP mRNA expression was detected in both regions following ic administration (with or without iv co-administration), while minimal mRNA was detected in brain following iv administration at P2 or P20 (Figure 1C).

At 3 weeks following neonatal injection (iv + ic route), immunostaining revealed presence of GFP protein in the liver and throughout the brain, including the cortex, hippocampus, and cerebellum (Figure 1D-G). Positive staining was detected in cells of differing morphology in the cortex suggesting transduction and promoter activity in both neurons and astrocytes.

### 3.2 AAV vector containing a mouse *Gldc* transgene reinstates mRNA and protein expression in GLDC-deficient mice

The AAV vector with CBh promoter was modified to contain the mouse *Gldc* (mGLDC) coding sequence encoding the consensus wild-type GLDC protein and packaged into AAV9 capsid. AAV9-CBh-mGLDC vector was administered to neonatal *Gldc^GT1/GT1^* mice by ic or iv + ic route. In the brain and liver of mice at 4 weeks post-administration, expression of Gldc mRNA and protein was analysed by quantitative real time RT-PCR (qRT-PCR) and immunoassay, respectively. *Gldc* mRNA was significantly more abundant in brain and liver of vector-treated mice than in vehicle-treated *Gldc^GT1/GT1^* mice (Figure 2A-B), with approximately 50-fold and 15-fold increase in expression in brain and liver, respectively. Similarly, GLDC protein was significantly more abundant in brain and liver of vector-treated *Gldc^GT1/GT1^* mice than in controls (Figure 2C-D). The fold-change increase in protein abundance was lower than for mRNA in the equivalent treatment groups. Expression of *Gldc* mRNA and protein in treated *Gldc^GT1/GT1^* mice typically exceeded endogenous wild-type levels in the brain (Figure 2A, C), with lower levels than wild-type in the liver (Figure 2B, D).

**Figure 2.**
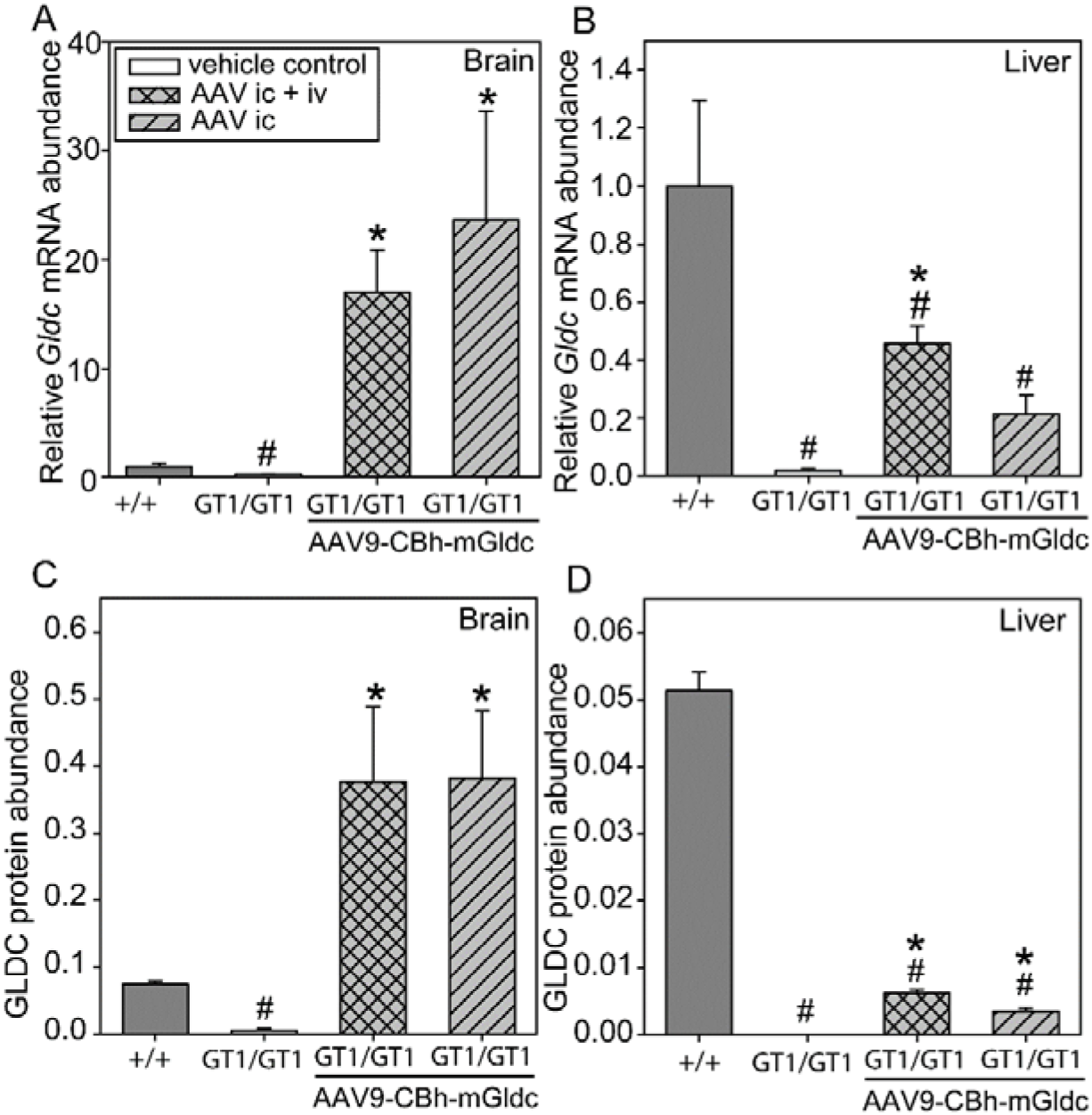
Expression of *Gldc* mRNA and protein following administration of AAV vector with mouse *Gldc* to neonatal *Gldc-*deficient mice. Abundance of *Gldc* mRNA (A-B) and GLDC protein (C-D) is lower in (A, C) brain and (B, D) liver of *Gldc-*deficient (*Gldc^GT1/GT1^*) mice compared with wild-type (+/+) at 6 weeks of age (^#^ significantly lower than +/+; p<0.001, t-test). Administration of AAV9-CBh-mGldc led to increase in mRNA and protein expression in (A, C) brain of *Gldc^GT1/GT1^*, by either iv +ic or ic route (* significantly different to untreated *Gldc^GT1/GT1^* or +/+, p<0.001; ANOVA), and (B, D) liver *Gldc ^GT1/GT1^*, by iv +ic route but not ic only (* significantly different to untreated *Gldc^GT1/GT1^* p<0.001; ANOVA, post-hoc analysis by Holm-Sidak method). In liver, mRNA and protein expression in treated *Gldc^GT1/GT1^* did not reach wild-type levels at 6 weeks of age (^#^ significantly lower than +/+; p<0.001, t-test). Number of samples (A-B) n = 6 +/+, 19 *Gldc^GT1/GT1^*, 9 iv + ic, 6 ic only brain samples; n = 5 +/+, 13 *Gldc^GT1/GT1^*, 13 iv + ic, 13 ic only liver. Number of samples for immunoassay was 5-7 per group.

Immunostaining for GLDC protein was used to confirm translation of the vector-expressed *Gldc* and assess distribution in the brain (Figure. 3). vector treated *Gldc^GT1/GT1^* mice showed widespread expression in of GLDC in the brain, including in hippocampus, cortex and cerebellum (Figure 3 D-F), unlike in control *Gldc^GT1/GT1^* mice, where GLDC protein was virtually undetectable (Figure 3 G-I). GLDC staining in vector-treated mice was of similar intensity to wild-type mice in hippocampus and cerebellum, and appeared more intense in the cortex. Hence, vector-mediated *Gldc* expression results in restoration of GLDC protein in the brain of treated mice, for at least 6 weeks after administration.

**Figure 3.**
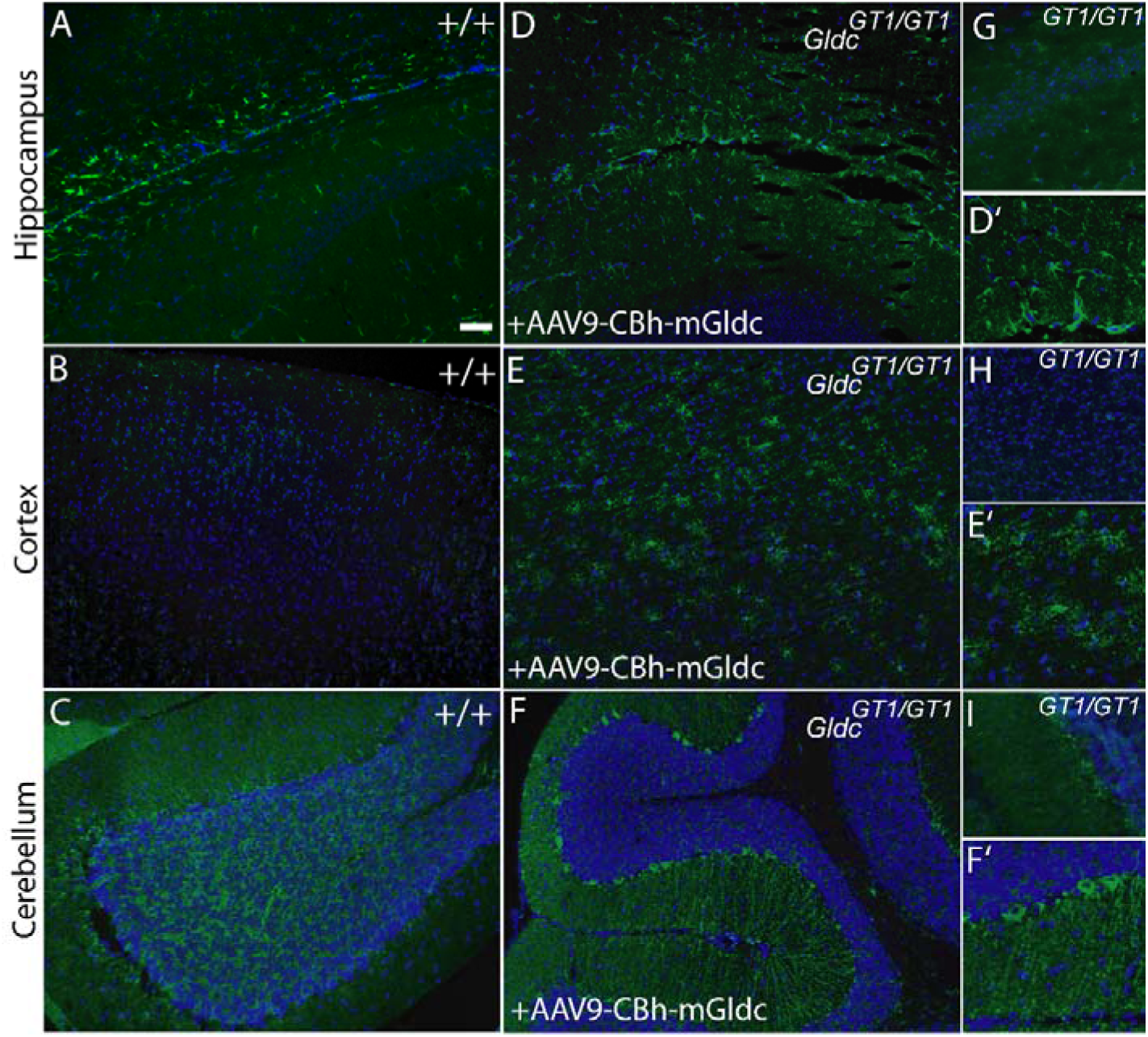
Expression of GLDC following administration of AAV vector expressing mouse *Gldc* to neonatal *Gldc-*deficient mice. Sites of GLDC expression in brain of +/+ mice include (A) hippocampus, (B) cortex, and (C) cerebellum. (D-F) After ic administration of AAV9-CBh-mGLDC at P1, GLDC protein is detected in brain of *Gldc^GT1/GT1^* mice after 6 weeks, but (G-I) is not is undetectable in untreated *Gldc^GT1/GT1^* mice. (D’-F’) show higher magnification images of F-H). Scale bar represents 100 µm (A-F are at same magnification).

### 3.3 Normalisation of metabolic biomarkers in *Gldc-*deficient mice treated with AAV9-CBh-mGldc

Elevated plasma glycine, the hallmark biochemical biomarker of NKH is recapitulated in the *Gldc-*deficient mouse model, and can be normalised by conditional *Gldc* expression in the liver [35]. At 6 weeks of age, plasma glycine concentration was significantly higher in *Gldc^GT1/GT1^* mice than +/+ littermates (Figure 3A), with a non-significant trend towards higher glycine in females than males among *Gldc^GT1/GT1^* mice at this stage. Among mice treated neonatally with iv AAV9-CBh-mGLDC, which leads to *Gldc* expression in liver, we observed significantly lower plasma glycine concentration than in untreated *Gldc^GT1/GT1^* mice (Figure 4A). As plasma glycine is regulated by liver GCS activity, this provides a measure of GLDC functionality. The treatment effect was observed in the overall cohort of mice and within male and female sub-groups. We conclude that vector expressed GLDC protein is functional and restores GCS activity.

**Figure 4.**
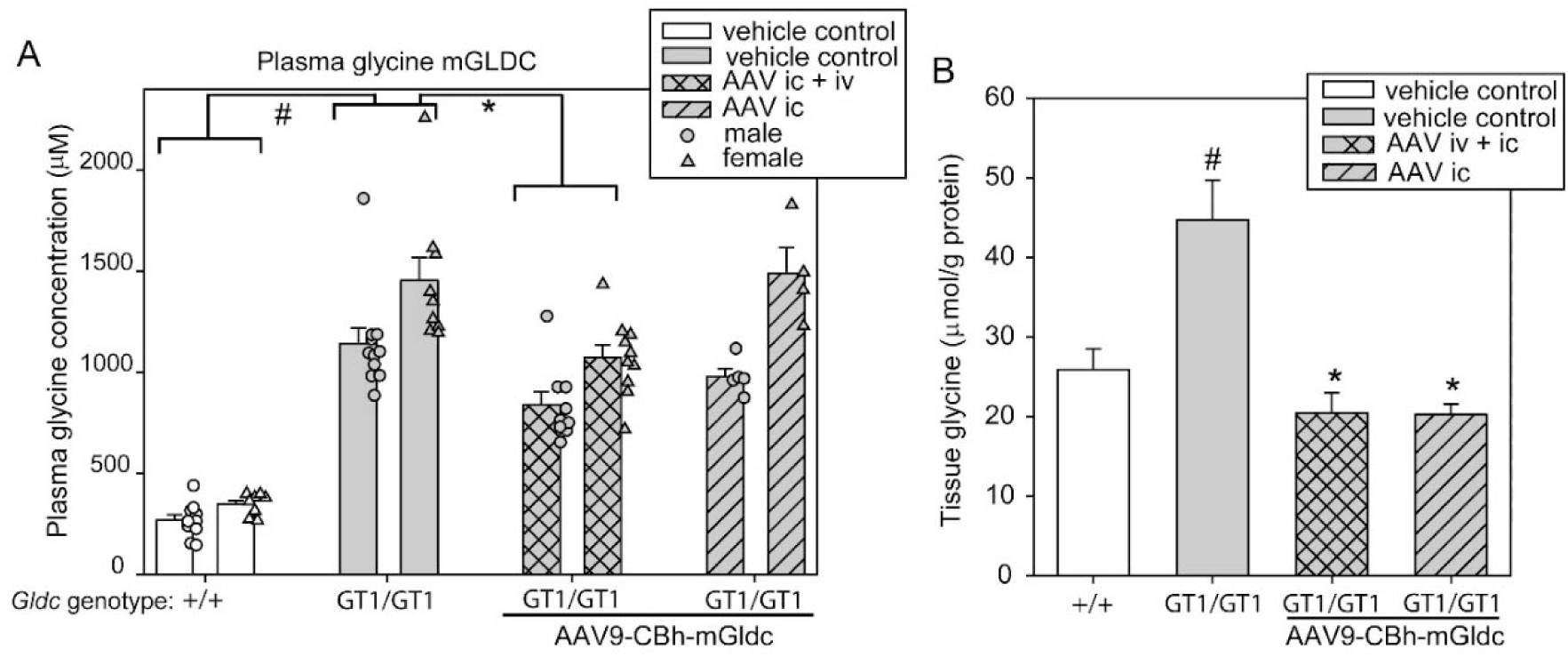
Administration of AAV-CBh-mGldc leads to lowering of plasma glycine. (A) Plasma glycine is elevated in vehicle treated *Gldc^GT1/GT1^* mice compared with wild-type (+/+) littermates (^#^significantly different, p<0.001). At 6 weeks after treatment at P1 plasma glycine was lower in *Gldc^GT1/GT1^* mice treated by iv+ic route, than in vehicle treated *Gldc^GT1/GT1^* mice (* significant difference, p<0.01). The effect of iv +ic treatment among *Gldc^GT1/GT1^* mice was observed among male and female mice analysed separately (significant difference between mice of same sex, p<0.05). (n = 19 +/+, 21 *Gldc^GT1/GT1^*, 19 *Gldc^GT1/GT1^* iv+ic treated, 9 *Gldc^GT1/GT1^* ic treated). (B) Brain tissue glycine content (normalised to protein) significantly varied between groups (p<0.001 ANOVA), but not between sexes (hence presented together). Glycine was significantly elevated in *Gldc^GT1/GT1^* compared with +/+ mice (^#^p<0.01) and was significantly lowered by AAV-CBh-mGldc treatment (*p<0.02). (n = 12 +/+, 16 *Gldc^GT1/GT1^*, 6 *Gldc^GT1/GT1^* iv+ic treated, 4 *Gldc^GT1/GT1^* ic treated).

As expected, we did not observe an effect on plasma glycine among mice in which *Gldc* expression was restored in brain but not liver, following vector administration by ic route only. However, we tested whether vector mediated GLDC expression in the brain was sufficient to lower tissue glycine content in *Gldc^GT1/GT1^* mice. We observed a significant reduction in brain tissue glycine content among mice 6 weeks after neonatal treatment with AAV9-CBh-mGLDC by iv + ic or by ic route only (Figure 4B). It therefore appears that excess glycine in brain tissue can be removed by GCS activity within the brain, even when plasma glycine is not fully normalised.

In parallel with evaluating glycine control, we tested whether treatment affected the supply of one-carbon units into FOCM. As reported at embryonic stages [34], we found that the folate profile is abnormal in *Gldc^GT1/GT1^* brain tissue, with an increase in relative abundance of THF and decrease in abundance of 5-methyl THF (Figure 5A-B). Following treatment with AAV9-CBh-mGLDC at P1, the folate profile in brain tissue of *Gldc^GT1/GT1^* mice at 6 weeks was normalised, with the relative proportions of THF and 5-methyl THF showing no difference to wild-type tissue (Figure 5A-B). As a further read-out of FOCM function in brain tissue we analysed abundance of betaine and choline. Choline is the precursor of betaine, which in turn can act as a methyl donor in the remethylation of homocysteine. We found that both metabolites are present at significantly lower levels in the brain of *Gldc^GT1/GT1^* mice (Figure 5C-D). Concomitant with normalisation of the folate profile we observed restoration of betaine and choline abundance following AAV-mediated restoration of *Gldc* expression by neonatal iv + ic or ic routes of administration (Figure 5C-D).

**Figure 5.**
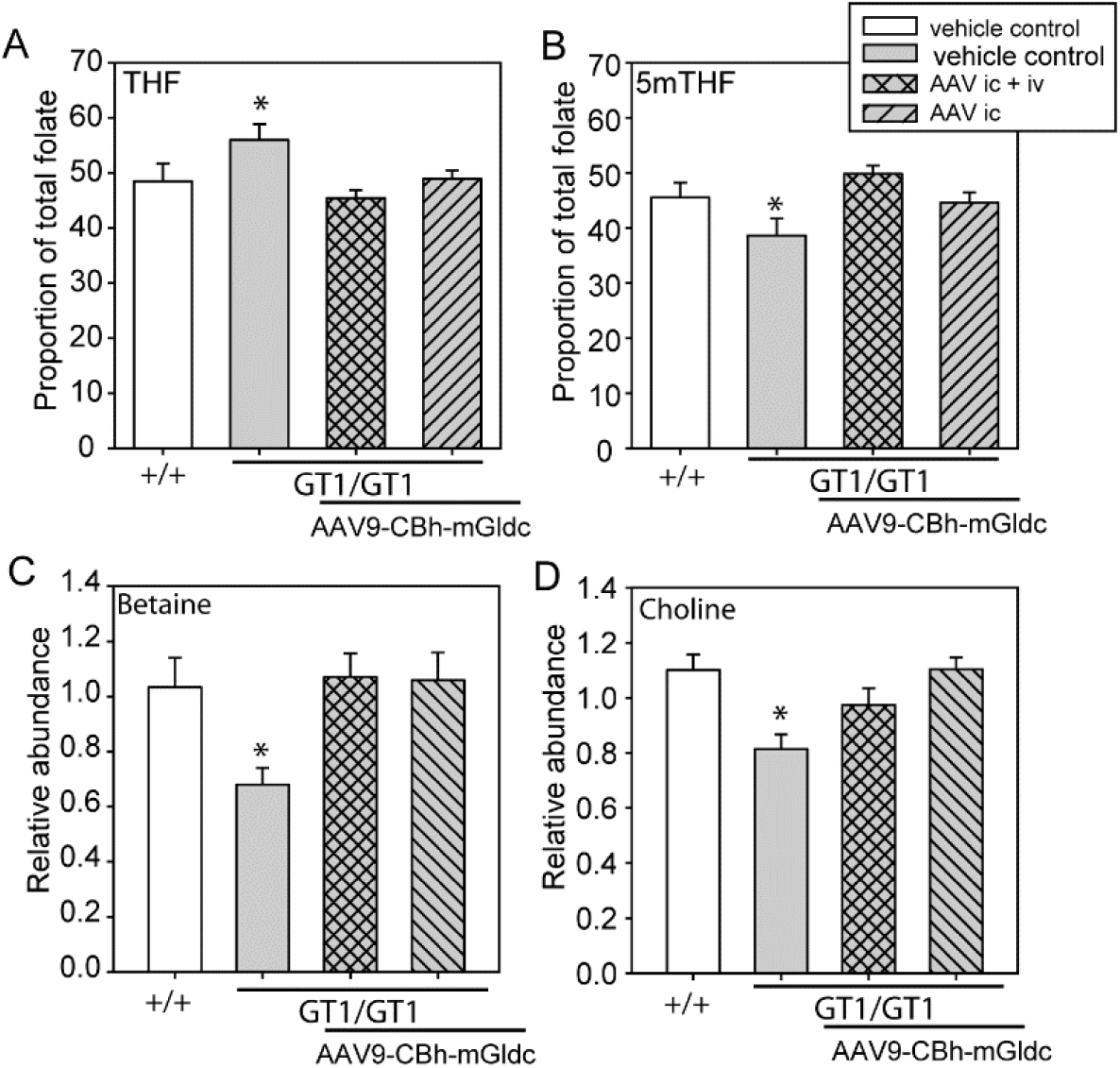
AAV-mediated GLDC expression normalises folate profile in brain of *Gldc-*deficient mice. (A-B) The abundance of (A) THF and (B) 5-methylTHF as a proportion of total folate differs in the brain of wild-type (+/+) and *Gldc-*deficient mice at 6 weeks of age (*significant difference between +/+ and *Gldc^GT1/GT1^* mice, p<0.05). No significant difference was observed following vector administration at P1 by iv + ic or ic routes (n = 5-6 mice per group). (C-D) Betaine and choline have significantly lower abundance in brain of *Gldc-*deficient mice than in wild-types at 6 weeks of age (*p<0.05; ANOVA), but abundance is normalised by administration of AAV-mGLDC by iv+ic or ic route at P1 (n = 6 mice per group).

### 3.4 AAV-mediated expression of functional human GLDC in mouse

Implementation of an AAV-based gene therapy for NKH will require a vector that leads to expression of the human GLDC protein. The GLDC protein in mouse (Uniprot Q91W43) and human (Uniprot P23378) comprise 1025 and 1020 amino acids respectively, with 92% identity. We hypothesised that the human protein would substitute for mouse GLDC in the GCS. Therefore, in order to test vectors expressing human GLDC we replaced the mouse *Gldc* coding sequence with a transgene sequence encoding the consensus human GLDC protein and packaged this into AAV9 capsid. The AAV9-CBh-hGLDC vector was administered to neonatal *Gldc^GT1/GT1^* mice and tissues were collected for analysis at 6 weeks of age. Expression of *GLDC* was confirmed in liver and brain by qRT-PCR. GLDC protein expression was assessed by immunoblot and as expected, endogenous mouse GLDC was detected in wild-type liver and brain but not in *Gldc^GT1/GT1^* mice (Figure 6A-B). In contrast, in samples from *Gldc^GT1/GT1^* mice that received neonatal treatment with AAV9-CBh-hGLDC, a GLDC band was present corresponding to vector-expressed GLDC. GLDC abundance in liver appeared to be lower than endogenous protein in liver but of greater abundance than endogenous protein in brain.

**Figure 6.**
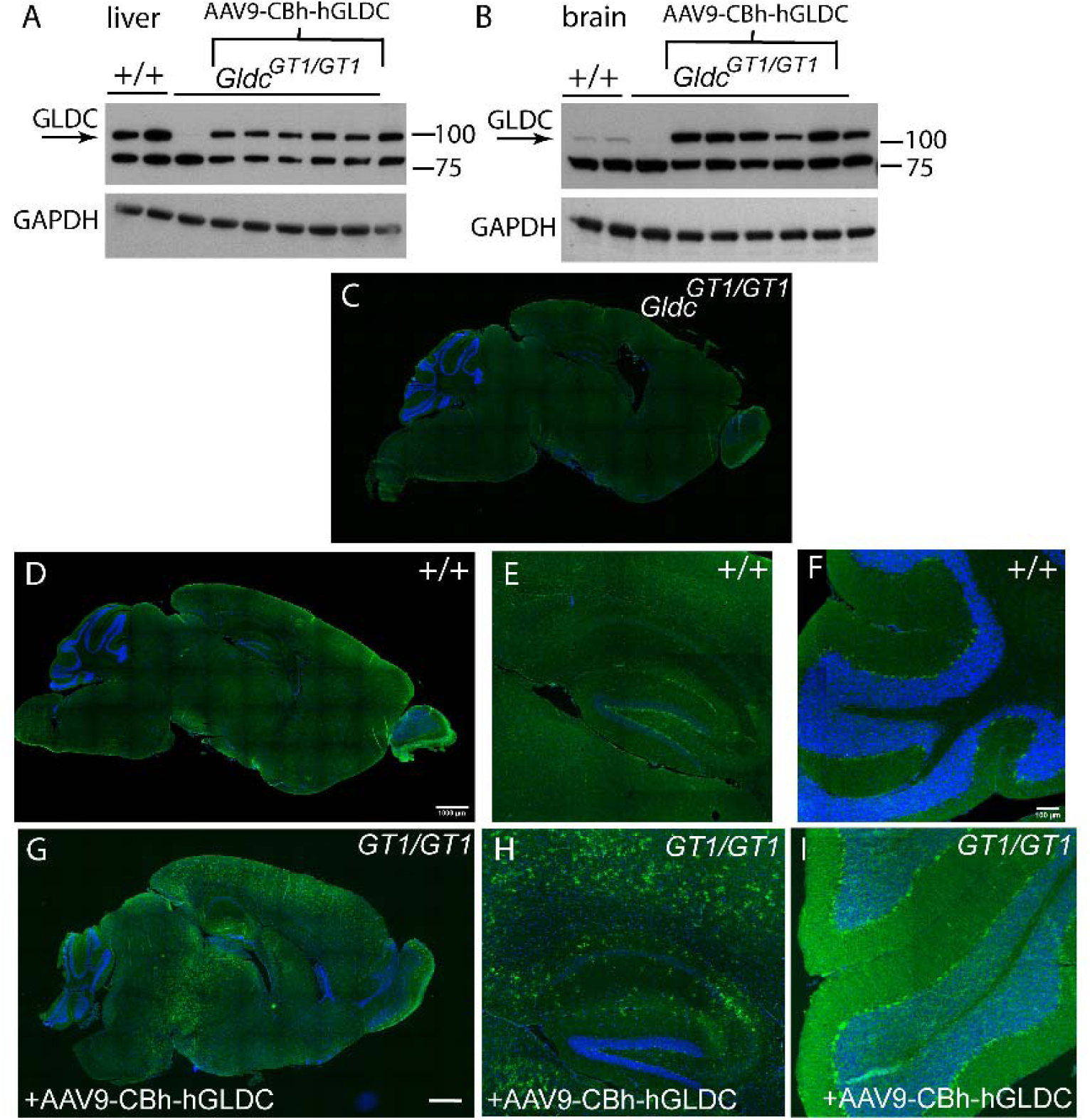
Expression of human GLDC protein following AAV vector administration to *Gldc-*deficient mice. (A-B) Immunoblot shows presence of endogenous mouse GLDC in (A) liver and (B) brain of wild-type (+/+) mice, but not in *Gldc-*deficient (*Gldc^GT1/GT1^*) mice at 6 weeks of age. Following administration of AAV9-CBh-hGLDC vector at P1 (iv + ic route), human GLDC protein is detected in both liver and brain of *Gldc^GT1/GT1^* mice. (C—I) Immunostaining for GLDC on sagittal sections of brain at 6 weeks of age reveals presence of GLDC protein in brain of (D-F) +/+ and (G-I) treated *Gldc^GT1/GT1^*, including hippocampus (E, H) and cerebellum (F, I), but not in untreated *Gldc^GT1/GT1^* mice (C). Whole brain images show tile scans of entire section (scale bar represents 1 mm).

We further examined expression of human GLDC in the brain of treated *Gldc-*deficient mice by immunostaining of sagittal sections. Unlike in untreated *Gldc^GT1/GT1^* brain (Figure 6C), GLDC staining was detected and appeared widespread in brain of AAV9-CBh-hGLDC treated *Gldc^GT1/GT1^* mice (Figure 6D-I), with intensity comparable to that of endogenous GLDC in untreated wild-type mice (Figure 6D-F).

Having confirmed that human GLDC is expressed in mice following AAV-vector administration we tested whether the human protein is functional in the context of the mouse GCS by analysis of plasma and brain tissue glycine (Figure 7). We observed significant lowering of plasma glycine at 6 weeks of age in vector-treated mice compared with untreated *Gldc^GT1/GT1^* mice (Figure 7A). In the brain of treated *Gldc^GT1/GT1^* the tissue glycine content was also significantly lowered compared to untreated *Gldc-*deficient mice, to a level that was comparable to wild-type (Figure 7B). Hence, vector expressed human GLDC is functional in the mouse.

**Figure 7.**
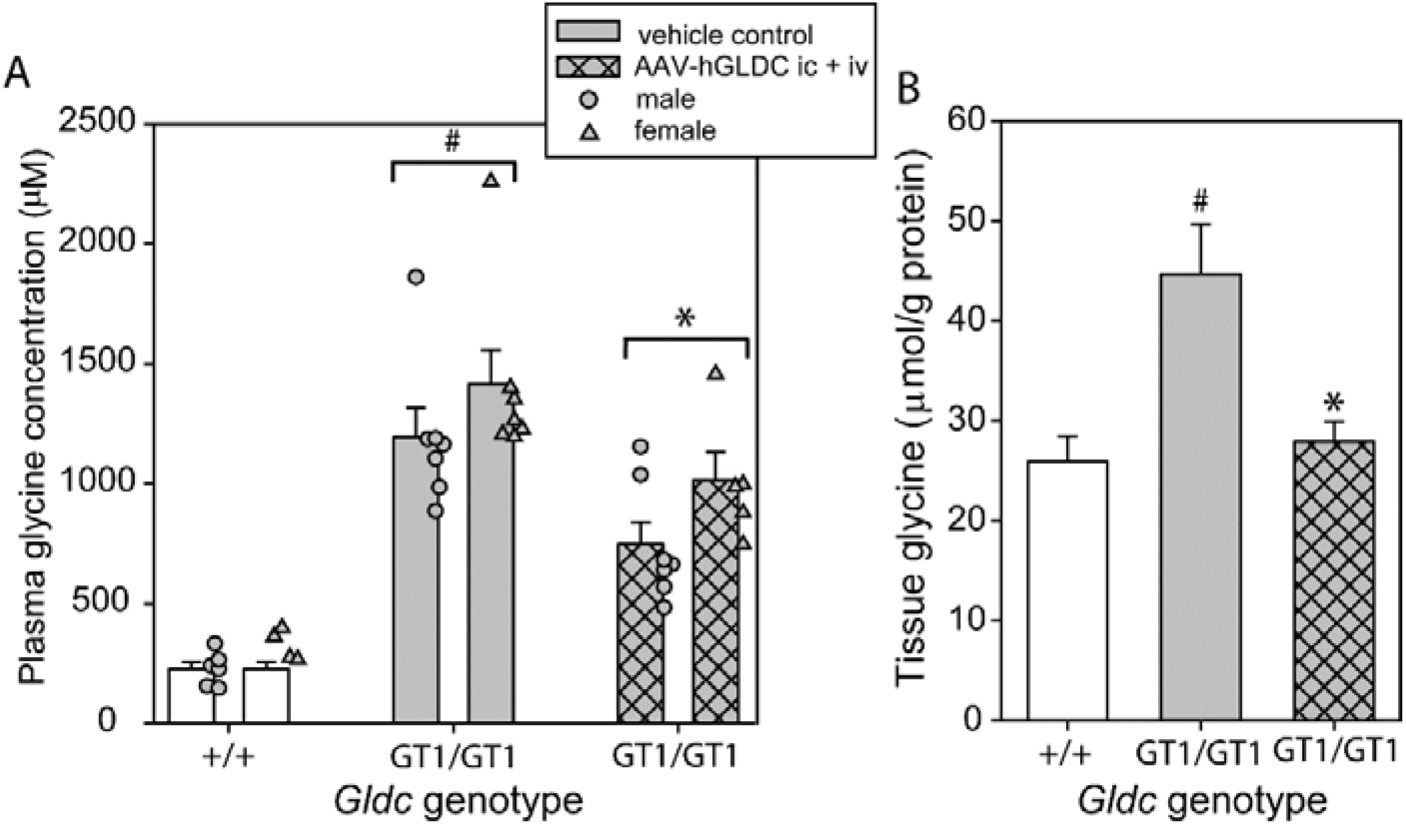
Lowering of plasma and brain tissue glycine by AAV vector mediated expression of human GLDC in *Gldc-*deficient mice. Plasma glycine concentration (A) and (B) brain tissue glycine (normalised to total protein content), are significantly elevated in *Gldc^GT1/GT1^* mice compared with +/+ at 6 weeks of age (^#^ p<0.001). Administration of AAV9-CBh-GLDC to neonatal mice by iv + ic route, leads to (A) significant lowering of plasma glycine compared with control *Gldc^GT1/GT1^* mice (* p<0.001 for overall group, and p<0.05 for individual sex comparisons), and (B) a significant reduction in brain tissue glycine abundance (* significant difference to *Gldc^GT1/GT1^* control group). Number of samples, n = 10-16 per group. Note that dataset for control +/+ and *Gldc^GT1/GT1^* overlaps with data in Figure 3 as AAV groups were treated in parallel.

## 4. Discussion

The GCS has two key functions, regulation of glycine concentration in tissues and supply of glycine-derived one carbon units to FOCM and downstream biochemical pathways. Disruption of each of these functions is thought to contribute to pathogenesis of NKH. As in humans, GCS disruption in mice leads to accumulation of glycine, and a series of glycine derivatives which may themselves be deleterious [34,35]. In parallel, impaired supply of one-carbon units leads to altered folate profile which we previously observed at embryonic stages and here confirmed in post-natal tissue. In addition to a shift in the folate profile, we found a reduced abundance of betaine and choline in the brain of *Gldc-*deficient mice which we hypothesise to reflect an increased requirement for betaine as a methyl donor owing to impaired supply of glycine-derived one-carbon groups via the GCS.

In the *Gldc-*deficient mouse model of NKH, treatment with benzoate or cinnamate is sufficient to lower circulating glycine [34,35], recapitulating the effect of benzoate in NKH patients [35]. However, this does not overcome suppression of FOCM and, as a therapeutic intervention in NKH, benzoate is associated with significant side effects including gastrointestinal irritation. Conversely, maternal supplementation of GLDC-deficient mice with formate, as a one-carbon donor, prevents the embryonic structural malformations that lead to neural tube defects and ventriculomegaly [34,35],[45], but does not lower excess glycine [34].

The ability to reinstate GCS function through vector-mediated expression of a functional copy of the mutated protein in NKH (GLDC or AMT) would provide an opportunity to restore both glycine regulation and one-carbon metabolism. In the current study we tested this concept in the GLDC*-*deficient mouse model, and found that AAV-mediated expression of GLDC resulted in both lowering of plasma and tissue glycine, and normalisation of FOCM intermediates. Lowering of plasma glycine was achieved following iv administration of vector, with a lesser effect following ic administration, suggesting that this effect largely reflects GLDC expression in the liver, the major site of GCS activity. However, within the brain the tissue glycine content was normalised by ic (or ic + iv) treatment indicating that GCS activity within the brain may be sufficient to regulate glycine within this tissue (even if not normalised systemically). Similarly, FOCM-related metabolites were restored by vector administration to the brain. This treatment led to widespread expression of GFP reporter or GLDC, including sites of endogenous expression in the hippocampus and cerebellum.

Expression of GLDC and normalisation of metabolite biomarkers in the brain was observed in young adult mice after neonatal administration, suggesting the potential for long-term benefit which will be evaluated in future studies. It was predicted that, as a predominantly non-integrating vector, the AAV vector may become ‘diluted’ relative to the total size of the liver owing to continuing cellular proliferation after neonatal administration of vector. However, significant lowering of plasma glycine, which reflects liver GCS activity, was still observed in young adults. Further studies will assess whether this effect persists longer-term. As a step towards long-term studies we found that neonatal administration of vector did not adversely affect survival or weight in wild-type mice maintained for up to 13 months.

Given that glycine conjugation and glycine cleavage are mediated by two separate enzyme systems, it is reasonable to predict that combined use of benzoate or cinnamate (to stimulate glycine conjugation [35]), with administration of a GLDC-expressing AAV vector (to reinstate GCS activity), could offer potential both for lowering plasma glycine and control of glycine, with restoration of FOCM within the brain, or other target tissues.

## Author contributions

**Kit-Yi Leung:** Conceptualisation, investigation, methodology, writing – original draft **Chloe Santos:** Investigation, visualisation **Sandra De Castro:** Investigation, methodology **Diana Gold Diaz:** Investigation **Andrew J. Copp:** writing – reviewing and editing, funding acquisition **Simon Waddington:** Conceptualisation, investigation, funding acquisition **Nicholas Greene:** Conceptualization, formal analysis, writing – original draft, reviewing and editing, supervision, funding acquisition, project administration

## Acknowledgments

This study was funded by LifeArc, Joseph’s Goal and Action Medical Research (grant number GN2403). We are also grateful for support from the Mikaere Foundation, Isla Rose Foundation and Miichen e.V. The authors thank Kyle O’Sullivan for technical assistance and Rajvinder Karda, Joanna Davidge, Stephanie Schorge, Paul Gissen, John Counsell and Julian Baruteau for helpful discussions. Research infrastructure and mass spectrometry at UCL GOS Institute Research was supported by the National Institute for Health Research Biomedical Research Centre at Great Ormond Street Hospital for Children NHS Foundation Trust and University College London.

## Notes

### Competing Interest Statement

The authors have declared no competing interest.

### Summary of Updates

Title has been modified. Minor edits to text and to acknowledgments

